# Long-term reciprocal gene flow in wild and domestic geese reveals complex domestication history

**DOI:** 10.1101/706226

**Authors:** Marja E. Heikkinen, Minna Ruokonen, Thomas A. White, Michelle M. Alexander, İslam Gündüz, Keith M. Dobney, Jouni Aspi, Jeremy B. Searle, Tanja Pyhäjärvi

## Abstract

Hybridization has frequently been observed between wild and domestic species and can substantially impact genetic diversity of both counterparts. Geese show some of the highest levels of interspecific hybridization across all bird orders, and two of the goose species in genus *Anser* have been domesticated providing excellent opportunity for joint study of domestication and hybridization. Until now, knowledge on the details of the goose domestication process has come from archaeological findings and historical writings supplemented with few studies based on mitochondrial DNA. Here, we used genome-wide markers to make the first genome-based inference of the timing of European goose domestication. We also analyzed the impact of hybridization on the genome-wide genetic variation in current populations of the European domestic goose and its wild progenitor: the greylag goose (*Anser anser*). Our dataset consisted of 58 wild greylags sampled around Eurasia and 75 domestic geese representing 14 breeds genotyped for 33,527 single nucleotide polymorphisms. Demographic reconstruction and clustering analysis suggested that divergence between wild and domestic geese around 5,300 generations ago was followed by long-term genetic exchange, and that greylag populations have 3.2–58.0% admixture proportions with domestic geese, with distinct geographic patterns. Surprisingly, many modern European breeds share considerable (> 10%) ancestry with Chinese domestic geese that is derived from the swan goose *Anser cygnoid*. We show that domestication process can progress despite continued and pervasive gene flow from the wild form.

**Significance Statement:** Reproductive isolation between conspecific wild and domestic populations is a cornerstone of the domestication process, yet gene flow between such wild and domestic populations has been frequently documented. European domestic geese and their wild progenitor (greylags) co-occur and can hybridize and we show that they represent a particularly persuasive case where wild and domestic populations are not isolated gene pools. Our study makes a first genome-based estimate of goose domestication, which up to now has mostly relied on archaeological findings and historical writings. We show ongoing gene flow between greylags and European domestic geese following domestication, but we also observe a surprisingly large contribution of Chinese domestic geese (a separate species) to the genetic make-up of European domestic geese.

## 1. Introduction

Reproductive isolation is a defining feature of speciation and yet hybridization between species is an important general phenomenon in evolution (1, 2). Among birds, the Anseriformes (ducks, geese and swans) show particularly pervasive hybridization, 41.6% to > 60% of species hybridizing with each other (3, 4). Domestication generates differentiated gene pools and reproductive isolation between domestics and their wild progenitor, but hybridization between domestic and wild forms has been well demonstrated in both plants (1, 5) and animals (6, 7). The impacts include genetic and trait enrichment of domestics, for instance, in chicken the acquisition of a yellow skin phenotype is a result of past mating between red junglefowl and grey junglefowl (8). In geese, the high tendency for hybridization among wild species is also shown among wild and domestic forms (9, 10), creating an exciting opportunity to study the complex dynamics of hybridization and domestication.

The domestic geese of the world (European and Chinese forms) are derived from two different wild species: the greylag (*Anser anser*) and the swan goose (*Anser cygnoid*) (11, 12). *A. anser* and *A. cygnoid* shared a common ancestor about 3.4 Mya (13), but are still able to hybridize (4), and some domestic breeds are reportedly hybrid (14). The greylag has been divided into the western, nominate subspecies *A. a. anser* (Linnaeus, 1758) with a European breeding range and the eastern subspecies *A. a. rubrirostris* (Swinhoe, 1871) breeding further east, although the subspecific boundary is not well defined, and mitochondrial DNA has not been found to distinguish them (10). Of these subspecies, *rubrirostris* is larger and lighter colored than *anser* (15) and has a pink bill and cold pink legs in contrast to the orange bill and flesh-colored legs of *anser*, the bill color used as primary evidence in favor of the original domestication of *rubrirostris* (16). As with all domesticates, domestic geese varieties are morphologically more diverse than their wild counterparts, particularly in plumage and body size (14).

The current knowledge about goose domestication relies largely on ancient texts and archaeological evidence. Questions about where and when domestication took place, the genetic changes associated with it and the later history of domestic geese, however, remain largely unresolved (10). There are depictions from the New Kingdom of Egypt that suggest geese were already fully domesticated by the 18^th^ Dynasty (1450-1341 BCE). The earliest reliable reference to domestic geese in western Eurasia is Homer’s Odyssey (first half of 8^th^ century BCE) and geese were certainly well-established poultry by Roman times (17).

Genetic diversity in the mitochondrial DNA (mtDNA) of greylag and European domestic geese showed reduced diversity in the domestics (10) which may result from an early domestication bottleneck or, alternatively, later breed formation. There is a particular mitochondrial haplogroup common in the domestics (10), and archaeological domestic goose bones from the High Medieval (11^th^-13^th^ century CE) of Russia belonged to that haplogroup (18).

MtDNA relationships between extant Chinese and European domestic goose breeds confirm that the former, excluding one breed, have swan goose ancestry, whereas European domestic goose and the Chinese Yili breed have greylag ancestry (12, 19, 20). However, Chinese mtDNA haplotypes may occasionally occur in European domestics, and vice versa (10, 19).

Genomic diversity data across populations is powerful in hybridization inference, and has been applied, for instance, to New World cattle, which along with their taurine ancestry were shown to have a greater proportion of indicine ancestry than previously assumed (21). Interpretation is still challenging and it is best to infer jointly the genetic impact of initial domestication and subsequent hybridization of wild and domestic populations, as the latter can obscure domestic-wild genetic relationships and may also give a false impression of the number of times a species has been domesticated (22–24).

Here we investigate goose domestication history using genome-wide single nucleotide polymorphism (SNP) data from thousands of loci, obtained by genotyping-by-sequencing (GBS). We used 56 and 50 samples of greylag and domestic geese from a previous mtDNA study (10), together with 2 new Turkish greylag and 25 new domestic specimens. We studied the interplay between domestication and hybridization by addressing the following questions: i) what is the extent of genetic differentiation amongst wild and domestic geese? ii) what is the approximate time of domestication? and iii) what is the role of intra- and interspecific hybridization in goose domestication history and the current genetic makeup?

## 2. Materials and methods

### a) Sampling

The greylag wild-collected samples derive widely from Eurasia (Fig. 1, *SI Appendix*, Table S1). The European domestic goose samples represented 14 different breeds (*SI Appendix*, Table S1) and individuals that did not belong to any recognized breed or were presumed to be crosses between European and Chinese domestic geese. Some specimens were reported to be Chinese domestic geese. The domestic samples were obtained from local breeders in Denmark, Sweden and the UK and those from Turkey were collected directly by the authors.

**Fig. 1.**
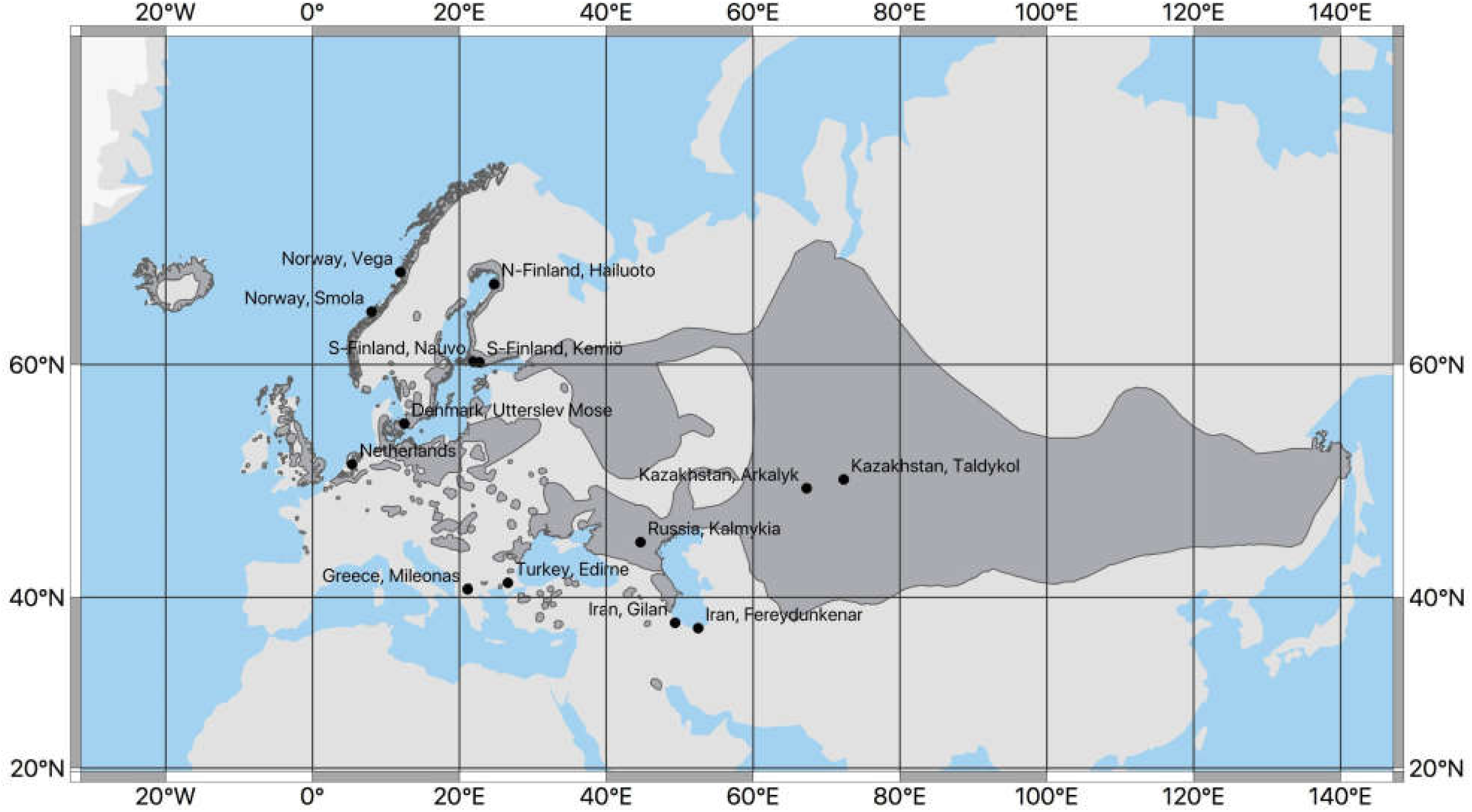
Map showing the sampling sites for wild greylags used in this study. The breeding area of the species is shown on darker grey. The sampling sites in Kazakhstan were combined for analyses (one sample per location) and the sampling sites in Southern Finland included combined samples from the geographically close sites of Västanfjärd, Nauvo (shown) and Kemiö (shown). The Iranian samples were collected during the wintering season. Map modified from IUCN (61).

### b) DNA extraction and GBS library construction

GBS (25) libraries were constructed at the Cornell Biotechnology Resource Center (BRC) following DNA extraction with the DNeasy Blood and Tissue Kit (QIAGEN) with RNase treatment. Each individual DNA sample and an adaptor with a unique barcode were combined in a 96-well plate along with a common adaptor. Samples were treated with the EcoT-22I (ATGCAT) restriction enzyme to create fragmented DNA. Barcoded adapters and common adapters with matching sticky ends were ligated to each sample with T4 DNA ligase. The samples were pooled and purified with a QIAquick PCR Purification Kit (QIAGEN). PCR amplification of the library used primers complementary to barcoded and common adapters with products purified as above, and the samples were 100 bp SE-sequenced with Illumina HiSeq 2000/2500 at the BRC.

### c) GBS pipeline and SNP calling

Raw sequence reads were run through the Command Line Interface of the Tassel 5 GBS v2 Discovery and Production pipelines (26). Details about the pipelines and SNP calling are in the *SI Appendix*. Good quality reads were recorded as tags and aligned to the *A. cygnoid domesticus* GenBank assembly (AnsCyg_PRJNA183603_v1.0 GCF_000971095.1) (27) using the Burrows-Wheeler Aligner with default settings (28). After running the raw data through the pipelines, 69,865 SNPs were obtained.

The SNPs were subjected to additional filtering using VCFtools (29). We removed indels, loci with more than two alleles and invariant loci. However, invariant loci were retained for phylogenetics, informing about greylag-swan goose divergence. After preliminary analyses loci with observed heterozygosity over 0.75 were removed as potential paralogs. Individuals with more than 20% missing data across loci were removed. The final dataset consisted of 33,527 biallelic SNPs and 133 individuals (58 wild and 75 domestic).

### d) Population structure analyses

Population clustering and structure was analyzed with STRUCTURE 2.3.4 (30) and Principal Component Analysis (PCA) (31). For the whole dataset, STRUCTURE was run with 1000 burn-in steps followed by 10 000 iterations of MCMC for data collection for *K* = 1-10 allowing admixture with five replicates of each run to reach convergence. For the STRUCTURE analyses done separately on greylags and European domestic geese, see *SI Appendix*. An admixture model with correlated allele frequencies among populations (32) was used in all STRUCTURE analyses and the iterations were automated with StrAuto 1.0 (33). We applied both likelihood of *K* and Δ*K* (34) of successive *K* values to determine the optimal number of clusters, using STRUCTURE HARVESTER (35). CLUMPP 1.1.2 (36) was used to align the assignments from different replicates of *K* and DISTRUCT 1.1 (37) for visualization. A PCA was performed with the prcomp function in R (38) and the significance of eigenvalues determined based on the Tracy-Widom distribution (22, 31).

A neighbor-joining tree was constructed for phylogenetic analysis, with pairwise distance between individuals obtained with the R package ape (39) based in 40,191 loci. The *A. cygnoid* reference genome and the invariant sites that differed from it were included in the tree construction.

### e) Tests for admixture and simulations of demographic history

The history of admixture was tested with a 3-Population test *f*_3_(C; A, B) implemented in AdmixTools 4.1 (40). This method offers a formal test to explain observed patterns of admixture in a target population without an outgroup. For identification of admixture between Chinese and European domestics, Grey and White Chinese were combined to represent the Chinese, and the Landes breed that had minimum indication of admixture in STRUCTURE was chosen to represent the European domestic source population. See also *SI Appendix* for further information

Different models of demographic history were tested with fastsimcoal2 ver 2.6 (41). We excluded all SNPs that had missing data within the whole data set and executed the analyses with a site frequency spectrum (SFS) based on 9,212 SNPs. As there are no estimates of the genetic diversity per base pair for greylags, we estimated the proportions of variable and monomorphic sites in the data. From the BAM file with –depth option in SAMtools 1.7 (42), we estimated 9,801,382 bp covered with GBS tags. We then mimicked the filtering steps done for the biallelic SNPs to reduce the total number of sites in equivalent proportions. We removed the same number of sites that corresponded to the number of SNPs that were removed because they were indels, had more than 2 alleles or had heterozygosity over 0.75. Since some of the SNPs were removed from this analysis due to missing data in some individuals, we removed an equal proportion of sites from the total number of sites as well. The final SFS had 1,681,316 sites.

To infer the demographic history, we chose a subset of individuals from both wild-collected greylags and domestic geese to represent the genetic variation in both groups. Therefore, 11 greylags with > 90.8% of greylag ancestry and 15 domestic geese with > 91.4% of European domestic goose ancestry were selected for the analysis. The mutation rate for the simulations was 1.38×10^−7^ (43). The parameter estimation for each model tested involved 100,000 simulations and 40 conditional maximization (ECM) cycles. The parameters for each model were estimated with 100 independent runs to obtain the global maximum. The models tested were i) simple divergence of two populations with no migration, ii) divergence of two populations with migration and iii) divergence of two populations with two different migration matrices (Fig. 2). The best model was selected based on Akaike’s weight of evidence as in (41). For parametric bootstrapping 100 SFS were simulated with the parameter estimates obtained from the real SFS, followed by maximum likelihood estimation with 50 independent runs for each bootstrap SFS. The 95% confidence intervals were obtained from the bootstrap data for each estimated parameter.

**Fig. 2.**
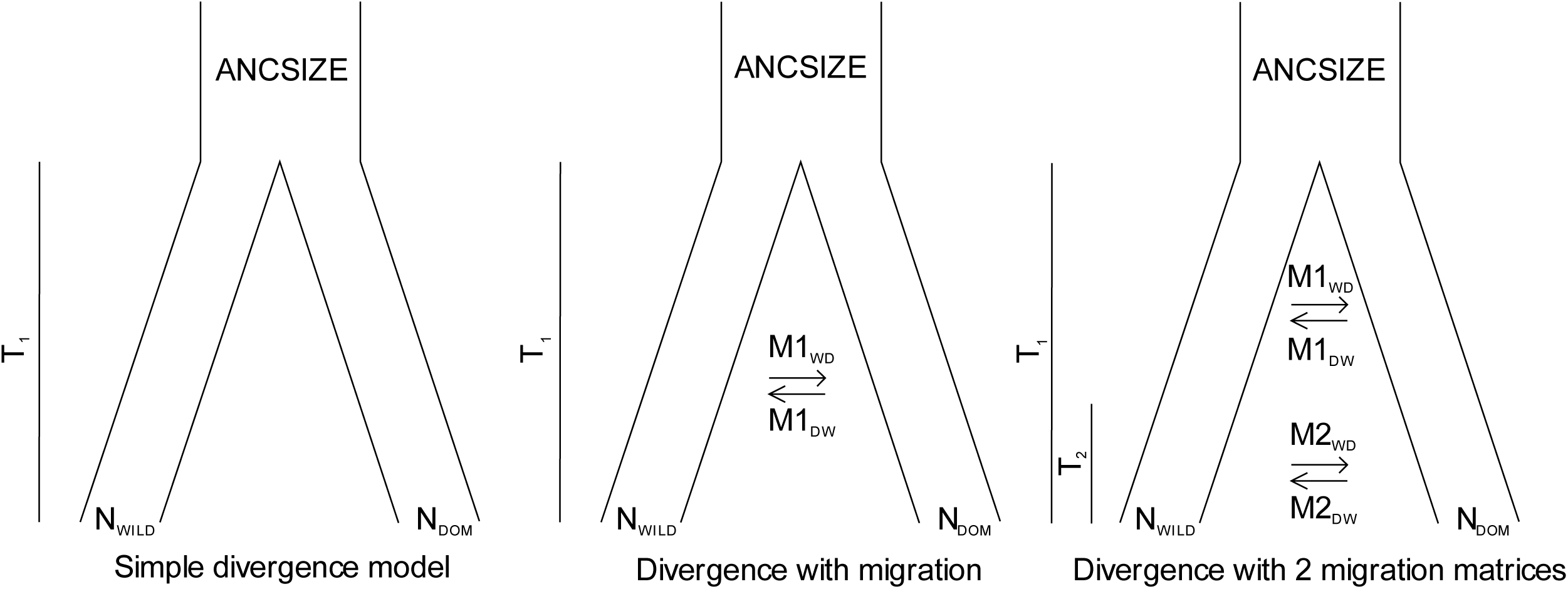
Demographic histories of goose domestication as tested with fastsimcoal2.

### f) The estimation of genetic diversity

Genetic diversity and pairwise *F*_*ST*_ values were investigated with the *hierfstat* R package (44). Expected heterozygosity (H_E_) was calculated for each locus and population and averaged across loci. Difference in average H_E_ between greylags and European domestics was tested with a two-sample t-test with the Welch correction for non-homogeneity of variance (45). For comparing the genetic diversity among wild and domestics, only pure greylag populations (< 10% admixture with domestic geese) and pure European domestic geese (< 10% admixture with Chinese domestic geese) were used to avoid hybridization effects on the estimates. The admixture proportions were obtained from STRUCTURE.

The variance components across loci for hierarchical F-statistics for pure greylags and pure European domestics were estimated using locus-by-locus analysis of molecular variance (AMOVA) implemented in Arlequin 3.5.2.1 (46). The significance was tested with 16 000 permutations.

## 3. Results

### a) Population structure

There was clear divergence between greylags and both European and Chinese domestic geese according to STRUCTURE and PCA (Fig. 3a-b). At *K* = 2, populations/breeds are clustered based on their status (wild or domestic) and, at *K* = 3, domestic geese are further separated into European and Chinese. At *K* = 4, the fourth cluster is admixed within greylag populations but none of the individuals are unanimously assigned to that cluster. The likelihood was highest for *K* = 3. These results were supported by PCA as the first two PCs out of 14 significant PCs (p < 0.05) were enough to separate the three groups (wild, European domestic, Chinese domestic) from each other (Fig. 3a). Overall the greylag populations showed 3.2% - 23.5% admixture proportions with European domestic geese when *K* = 3. In contrast, not all European domestic geese showed admixture with greylags and the admixture percentages ranged from 0.0 to 8.4%. At *K* = 3 many European domestic goose breeds showed mixed ancestry with Chinese domestic geese (0.0-27.1%).

**Fig. 3.**
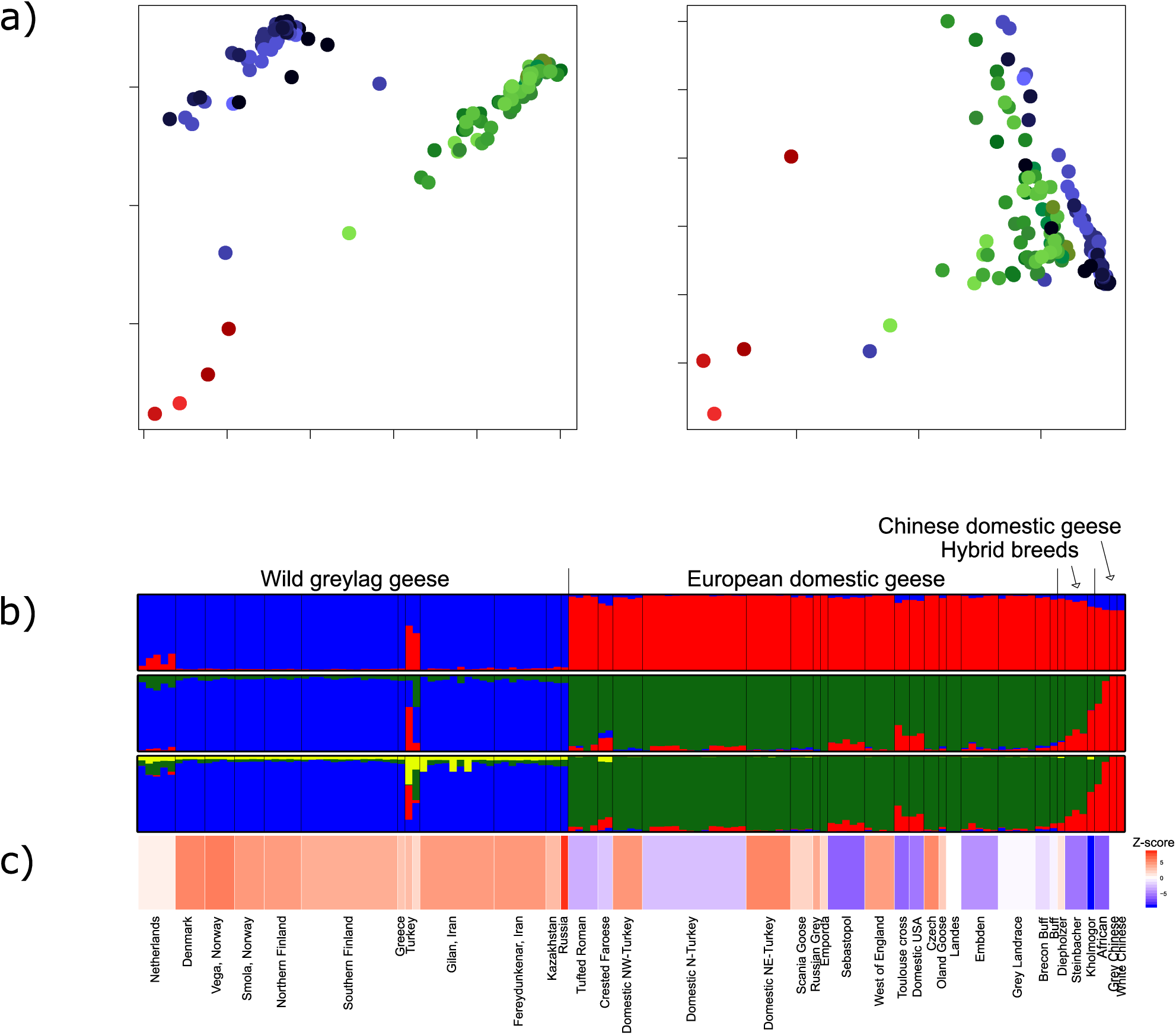
The genetic divergence and hybridization patterns in greylag and domestic geese. Population status and names labelled as in *SI Appendix*, Table S1. The colors in a) and b) are associated to different groups as follows: greylags (blue), European domestics (green) and Chinese domestics (red). a) The first three principal components summarizing the genetic variation in geese (percentage explained by each PC is shown). b) STRUCTURE assignment plots for *K* = 2, *K* = 3, and *K* = 4. Each vertical line represents one individual with *K* number of colors indicating proportion of ancestry from the inferred clusters. c) Plot relating to the *f*_3_ (*SI Appendix*, Table S2) values obtained for each population. The more negative the *f*_3_, the more significant is Z-score in favor of admixture. The *f*_3_ values were not calculated for Landes and the Chinese geese, as they were used as source populations, thus they were given an *f*_3_ value of 0.

The neighbor-joining tree repeated the major patterns observed with STRUCTURE and PCA revealing a star shaped phylogeny and confirming that the domestic and greylag geese largely form different clades (*SI Appendix*, Fig. S1). Surprisingly, the Chinese domestic geese were closer to European domestic geese and greylags, than to the swan goose reference genome. In addition, one greylag from Turkey was more closely related to the Chinese domestic geese than other greylags, also indicated by admixture proportions from STRUCTURE. Further, two Crested Faroese individuals and four domestics from the USA (2 unknown and 2 Toulouse crosses) were closer to Chinese than European domestic geese. These six individuals also showed high proportions of admixture with Chinese domestics in the STRUCTURE analysis.

Some further population structure was observed within both greylags and domestic geese, when analyzed separately with STRUCTURE and PCA. Geographically, greylags differentiated by subspecies (*SI Appendix*, Fig S2-S3). STRUCTURE indicated little differentiation among European domestic geese, but the PCA revealed separation between the European breeds and the Turkish domestic geese (*SI Appendix*, Fig S4-S5).

### b) Admixture and the time of domestication

Although, STRUCTURE implied considerable mixed ancestry from multiple genetic clusters for Dutch and Turkish greylags, the *f*_3_ analysis did not confirm admixture for these populations. The two Turkish greylag samples came from the same area as our NW-Turkish domestic population, which among Turkish domestic geese showed highest admixture with greylags (2.2%), but admixture was not confirmed with the *f*_3_ test.

The *f*_3_ analysis confirmed admixture of domestic geese in line with STRUCTURE results. Most notably, the African breed is a hybrid between European and Chinese domestic geese (Z-score −6.399), unexpected as this breed has been assumed to have originated solely from swan goose. The European-Chinese hybrid status of the Kholmogor and Steinbacher breeds was also confirmed (Z-scores of −8.933 and −5.349, respectively). The Kholmogor breed also fell halfway between European and domestic geese both in STRUCTURE and PCA, whereas the Steinbacher was genetically closer to European domestic geese in the PCA. However, the Diepholzer breed, which reportedly is also a hybrid, was not confirmed as such in our analysis. Other domestic breeds/groups with admixture status in STRUCTURE were also confirmed to have a European-Chinese admixture when a Z-score threshold of −3 (roughly corresponding to *p* < 0.01) was used: Sebastopol, Toulouse cross, Domestic NY, Embden, Tufted Roman (Fig. 3c, *SI Appendix*, Table S2). The Crested Faroese breed gave indication of admixture based on STRUCTURE analysis and the *f*_3_ test supported this (Z-score of −2.228, p < 0.05). Surprisingly, the Northern Turkish domestic population was not admixed with Chinese domestic geese in STRUCTURE but *f*_3_ analysis gave a contrasting signal (Z-score −2.459, p < 0.05).

The demographic model that best fit our data suggested divergence of greylag and domestic geese with a recent migration rate change (Table 1, *SI Appendix*, Table S3). The model suggested divergence around 5319 generations ago (95% confidence intervals (CI): 1538-8225) with asymmetric but close to equal migration rates from greylags to domestic geese following divergence. About 159 (79-462) generations ago, the migration matrix changed, suggesting higher gene flow from domestic geese to greylags towards modern times. Given an estimated generation time for these geese of about 3 years, this suggests divergence about 14 000 BCE and gene flow shift about 480 years ago.

**Table 1.** The parameter values that maximize likelihood estimates (MLE) of the preferred demographic model for goose domestication history (see text) with their 95% confidence intervals (CI) and the search range for the parameter values.

| Model | Number of parameters | Parameter | MLE | 95% CI | Search range |
| --- | --- | --- | --- | --- | --- |
| Divergence with 2 migration matrices | 9 | ANCSIZE | 1112 | 291.95 - 9030.75 | 100 - 100000 |
|  |  | T<sub>1</sub> | 5319 | 1538.80 - 8225.45 | 100 - 20000 |
|  |  | M1<sub>WD</sub> | 5.35x10<sup>-4</sup> | 2.96x10<sup>-4</sup> - 6.75x10<sup>-4</sup> | 1x10<sup>-5</sup> - 20 |
|  |  | M1<sub>DW</sub> | 4.25x10<sup>-4</sup> | 1.73x10<sup>-4</sup> - 6.03x10<sup>-4</sup> | 1x10<sup>-5</sup> - 20 |
|  |  | T<sub>2</sub> | 159 | 79.65 - 462.95 | 50 - 10000 |
|  |  | M2<sub>WD</sub> | 6.69x10<sup>-4</sup> | 4.57x10<sup>-4</sup> - 9.69x10<sup>-4</sup> | 1x10<sup>-5</sup> - 20 |
|  |  | M2<sub>DW</sub> | 1.72x10<sup>-3</sup> | 1.33x10<sup>-3</sup> - 2.15x10<sup>-3</sup> | 1x10<sup>-5</sup> - 20 |
|  |  | N<sub>DOM</sub> | 959 | 879.95 - 1073.25 | 50 - 100000 |
|  |  | N<sub>WILD</sub> | 2504 | 2341.60 - 2695.55 | 50 - 100000 |

ANCSIZE, effective population size of ancestral population; T<sub>1</sub>, time of divergence in generations; N<sub>DOM</sub>, effective population size for domestic geese; N<sub>WILD</sub>, effective population size for greylags; T<sub>2</sub>, estimate of time in generations when the migration matrix switched; M1<sub>WD</sub> migration rate from wild to domestic following T<sub>1</sub>; M1<sub>DW</sub> migration rate from domestic to wild following T<sub>1</sub>; M2<sub>WD</sub> migration rate from wild to domestic following T<sub>2</sub>; M2<sub>DW</sub> migration rate from domestic to wild following T<sub>2</sub>.

### c) Genetic diversity

An AMOVA was used to partition genetic diversity among greylag vs. domestic (group level), and among populations (greylag) and among breeds (domestics), and within population levels (Table 2). The fixation index between greylag and domestic geese was 0.158 and there was also significant differentiation among greylag populations/domestic breeds. The average pairwise F_ST_ between greylag populations and domestic breeds was 0.197, among greylag populations 0.088 and among domestic breeds 0.174 (*SI Appendix*, Table S4).

**Table 2.** Hierarchical analysis of molecular variance (AMOVA) of greylags and their domestic descendants, considering pure populations of greylags (first group) and pure breeds of European domestic geese (second group).

| Source of variation | Sum of squares | Variance components | Percentage variation | Fixation indices |
| --- | --- | --- | --- | --- |
| Among groups | 47565.119 | 431.4291 | 15.8 | $F_{CT} = 0.158^{**}$ |
| Among populations and breeds within groups | 82960.489 | 302.51404 | 11.1 | $F_{SC} = 0.131^{**}$ |
| Within populations and breeds | 345889.821 | 2003.45893 | 73.2 | $F_{ST} = 0.268^{**}$ |
| Total | 476415.429 | 2737.40207 |  |  |

\*\* p < 0.001

The genetic diversity measured as average H_E_ was higher in pure greylags (0.146) than pure European domestic geese (0.096) (Welch’s t-test, degrees of freedom (df) = 10.594, *p* = 3.91×10^−5^). The difference in average H_E_ remained when non-pure greylag and non-pure European domestics were included in the comparison (0.156 vs. 0.107; Welch’s t-test, df = 19.28, *p* = 0.000418) (*SI Appendix*, Table S1). The Steinbacher and African breeds (*SI Appendix*, Table S1) were excluded from both estimates because the Steinbacher is known to be a European-Chinese cross and the African has swan goose ancestry. The average H_E_ ranged from 0.140 (Denmark) to 0.150 (Kazakhstan) in pure greylags and from 0.047 (Landes) to 0.123 (Domestic N-Turkey) in pure European domestics. The average H_E_ was higher in greylag populations that showed high admixture proportions with domestic geese, Netherlands (0.157, 13.8% European domestic, 2.2% Chinese domestic) and Turkey (0.236, 23.5% European domestic, 34.5% Chinese domestic), but not significantly so (Welch’s t-test, df = 1.002, *p* = 0.421, average H_E_: pure greylags 0.146 vs non-pure greylags 0.196). The average H_E_ was also higher in European domestics that showed high admixture proportions with Chinese domestics, i.e. Crested Faroese (0.117, 17% Chinese domestic), Sebastopol (0.133, 11.1% Chinese domestic), Domestic NY (0.143, 78.6% European domestic, 21.4% Chinese domestic) and Toulouse cross (0.151, 72.9% European domestic, 27.1% Chinese domestic), and the difference was also statistically significant (Welch’s t-test, df = 9.0991, *p* = 0.0039, average H_E_: pure European domestics 0.096 vs non-pure European domestics 0.136).

## 4. Discussion

We studied the dynamics of domestication and hybridization in grey (*Anser*) geese using genome-wide SNP data. The results demonstrated genetic divergence between Eurasian wild greylag and European domestic geese with long-term genetic exchange between them. We also inferred temporal changes in the direction of gene flow. The degree of hybridization between greylag and domestic geese also varied geographically. Surprisingly, several domestic goose breeds also showed a substantial genetic contribution of Chinese domestic geese. We also provide insights about the origin and the timing of goose domestication.

### a) Genetic diversity and differentiation of greylag and European domestic geese

Domestic species often show reduced genetic diversity compared to their wild ancestor, attributable to genetic drift during population bottlenecks of initial domestication, combined with subsequent artificial selection associated with breed formation (47). Domestic geese appear to follow the same trend. We found European domestic geese to have lower H_E_ than wild greylags.

European domestic geese are genetically distinct from their wild progenitor but no more so than for other domestic birds. The average pairwise F_ST_ values between greylag populations and domestic goose breeds were lower than between red junglefowl and domestic chicken populations (48), and domestic geese are less distinctive than domestic pigeons (49). Among domestic geese, the Turkish are particularly interesting. From mtDNA, the Turkish domestic geese stand out as the most genetically variable group (10), and although this is less evident from GBS, among the pure European domestic geese the Northern Turkish showed the highest average H_E_. The *f*3 analysis indicates a history of admixture with Chinese domestics for this population, which may explain its high genetic diversity.

We found a genetic separation between European and Near Eastern populations of greylags that aligned with the western and eastern subspecies (*A. a. anser* and *A. a. rubrirostris*) (50), a distinction which could not be made based on mtDNA (10). Hybridization between the western and eastern subspecies is suggested from admixture in Dutch and Danish greylags in STRUCTURE. There is historical evidence for the introduction of *rubrirostris* to Belgium in 1954 and to Netherlands in 1960s (9, 51); thus, *rubrirostris* genes may have originated from the recently introduced gene pool spreading to Denmark.

### b) When and where were geese domesticated?

Traditional views on goose domestication claim it first occurred in the eastern Mediterranean (possibly Egypt) around the 3^rd^ Millennium BCE (17, 52). Domestication of chicken and perhaps pigeon took place earlier, but domestication of duck later, at least in Europe (23). Demographic modelling suggests that the wild greylag and related domestic lineages split approximately 5,300 generations ago placing domestication origins at 14 000 BCE assuming a 3-year generation time (15). This estimated genetic divergence time is, admittedly, considerably earlier than any evidence for animal domestication except dog. It is important to note that the estimated divergence times have large confidence intervals and merely indicates the split between the ancestors of contemporary wild and domestic lineages. It is most likely that our demographic modelling reflects the early divergence of different lineages of greylags, only one of which contributed to later domestication. The subsequent reduction or even disappearance of that wild lineage means that, despite wide geographical sampling, the possible modern wild population(s) of the greylag progenitor to domestic geese was not sampled in this study and must, therefore, be sought by ancient DNA approaches.

Given that genetic diversity would be expected to be highest in the ‘domestication center’ and reduce with increasing distance from there, the high mtDNA diversity of Turkish domestic geese means the eastern Mediterranean remains a strong candidate for the origin of goose domestication, and our nuclear data accord with that. However, as we have shown, hybridization between wild and domestic geese can also generate high genetic diversity both within and outside the original domestication location.

### c) The role of intra- and interspecific hybridization in goose domestication history

#### i. Evidence of current hybridization

Domestic animals and their wild relatives are often observed to interbreed, and this is also true for geese. Both field observations and mtDNA results (9, 10) suggested some current hybridization between domestic and greylag geese. Genome-wide analysis covering multiple greylag populations and domestic breeds revealed a considerable impact of hybridization on genetic diversity of both wild and domestic geese.

Hybridization is particularly prevalent in certain geographical regions. Dutch and especially Turkish wild greylag samples had more shared genetic affiliation with domestics than Scandinavian and Finnish greylag populations (Fig. 3b). Some regions may offer more hybridization opportunities, e.g. climate may allow greylags to be sedentary year-round and be favorable for keeping domestic geese. The Netherlands, for instance, lies on the Atlantic flyway offering breeding, staging and wintering areas for greylags (53, 54). Since pair-bonding of geese generally occurs on wintering grounds (55), hubs for migrating geese such as the Netherlands may permit population mingling. Nevertheless, the *f*_3_ test did not support a simple history of admixture for the Netherlands. Patterson et al. (40) have stated that population-specific drift may mask the signal of admixture in such analyses, leading to a non-negative *f*_3_. The *f*_3_ model is relatively simple, with only two sources, and may not catch the signal of admixture in the Dutch greylag population because of the previous contribution of *rubrirostris*, which was not included in the model.

Based on ringing data most greylag populations in Scandinavia follow the Atlantic flyway - some of the geese wintering in the Netherlands and others in southwest Spain. However, Finnish greylags favor the Central European flyway and winter in North Africa, with a minority of Finnish greylags using the Atlantic Flyway (53, 54). The Finnish populations of greylag showed the lowest admixture proportions with domestic geese (S-Finland 3.2% domestic goose, N-Finland 3.3% domestic geese) among greylag populations. Rearing geese is not a popular practice in Finland and they constituted less than 5% of poultry kept in Finland in 2014 (56). The Norwegian populations showed only slightly higher admixture proportions with domestic geese, although domestic mtDNA haplotype ANS19 was detected from a wild greylag collected in Finnmark, Norway (57). This haplotype is a partial sequence of the D5 haplotype identified by Heikkinen et al. (10), and identical to that found in White Roman domestic geese (58).

Inferring the hybridization patterns in the Turkish greylags is more complicated, as Turkish greylags indicate hybridization with both Chinese and European domestics. Both greylags sampled in Turkey showed considerable admixture with domestic geese. One of them appeared genetically as a hybrid of European and Chinese domestic goose with only a small proportion of greylag ancestry, whereas the other one was a more equal mix of European domestic goose and greylag supplemented by a considerable Chinese domestic goose ancestry. There is some indication of hybridization between greylags and domestic geese within that area as the domestic geese sampled from the same area showed some admixture with greylags, but this was not confirmed with *f*_3_ analysis. These results may reflect a local practice of keeping captive greylags within a flock of domestic geese. Another possibility is that the Turkish greylags have hybridized with some unsampled distinct greylag population and simply appear genetically like domestic geese due to lack of representation of the unsampled wild population. The greylag population breeding and wintering in the Black Sea region is not well monitored (59).

#### ii. Long-term hybridization

Domestication can be seen as an analogy of speciation where an animal population transforms to an ecotype that is adapted to the human niche (23) and at later stages of domestication is perpetuated with reproductive isolation in the form of selection managed by humans (60). However, this reproductive isolation may not be complete (7). While the genetic divergence of the greylag and its domestic descendant is evident, our results suggest extensive long-term genetic exchange between them. In addition, the demographic modelling suggests that the gene flow patterns have changed over time.

Initially, gene flow was greater from greylag to domestic geese. Possibly early goose herders supplemented the domestic gene pool with wild-caught greylags. Alternatively, wild ganders may have interbred with domestic geese and contributed to the domestic gene pool. It is unlikely that the early stages of goose domestication were rigorously managed, allowing matings outside the domestic gene pool. Both would explain the observed higher gene flow from greylag to domestic geese in the early stages of domestication. It is in the farmers’ interest to keep the domestic geese and wild geese reproductively isolated to keep control over the traits that are being selected, but artificial selection of traits would have become possible only after the domestic gene pool had been established. After that, it may occasionally be beneficial to restock the flock to maintain enough genetic diversity. The natural tendency for imprinting in geese facilitates this practice. Later, the balance of gene flow shifted towards greylags. This likely occurred after goose-keeping became well-established in the Medieval period (17). The rise in number of domestic geese may have allowed an increase in domestic goose escapees resulting in increased gene flow from domestic geese back to greylags.

Furthermore, not only have domestic geese admixed with wild greylags but also European and Chinese domestic geese have hybridized. Hybridization with ancestral species or closely related species is frequent in domestic species, e.g., the genetic makeup of chicken derives from multiple different species of *Gallus* (8). Similarly, the genetic make-up of domestic geese seems to derive from two closely related species. This hybridization with Chinese domestic geese may have introduced some traits not present in greylags to European domestic geese and vice versa.

## Authors’ contributions

MEH conceived the study, contributed to data collection, analyzed the data and drafted the manuscript. MR acquired the funding, conceived the study and contributed to data collection. TAW contributed to data collection and participated to data analysis. MMA and IG contributed to data collection. KMD, JA, JBS and TP conceived the study and contributed to writing and interpretation of data, with TP also participating in data analysis. All authors, excluding MR, reviewed, improved and approved the manuscript.

## Acknowledgements

We thank the Ministeriet for Fødevarer, Landbrug og Fiskeri in Denmark, the Svenska Lanthönsklubben in Sweden and the Goose Club in the UK for their help in obtaining the domestic goose samples. Silvia Markova is acknowledged for her help with the laboratory work. We are grateful for Tiina Mattila for her help with the fastsimcoal2 analysis. Lastly, we thank the CSC – IT Center for Science in Finland for providing computing resources.

## Extended methods

### GBS pipeline and SNP calling

Raw sequence reads from Illumina were run through the Command Line Interface of the Tassel 5 GBS v2 Discovery and Production pipelines (1) on the Taito supercluster maintained by the CSC - IT Center for Science in Finland. The GBSSeqToTagDBPlugin was run with the default setting except for the minimum quality score set to 20. This plugin identifies good quality reads with the barcode and cut site from the raw sequence data and trims off barcodes and truncates sequences if another cut site is found in the sequence. The reads that were pulled from the raw data were trimmed to 64 bp (base pairs) but if a second cut site was found in the read, the sequence was truncated and kept if the remaining sequence was longer than 20 bp. The good quality reads are recorded as tags and along with individuals in which they appear, the tags are stored in the local database. Given the parameters used, the length of each good quality tag ranged from 20 to 64 bp and each position in the sequence had a minimum quality score of 20 from Illumina sequencing. After this, the tags were aligned to the *A. cygnoid domesticus* GenBank assembly (AnsCyg_PRJNA183603_v1.0 GCF_000971095.1) (2) using the Burrows-Wheeler Aligner with default settings (3). Then, the SAMToGBSdbPlugin was run with default settings to determine the potential positions of tags in the reference genome and the position information was recorded to the local database. Altogether 285,760 tags were mapped on the reference genome and 66,163 tags were left unmapped. In the next step the DiscoverySNPCallerPluginV2 was used to align the tags positioned in the same physical location with each other and called single nucleotide differences between aligned tags as SNPs. The SNP position and allele data were stored in the local database. The DiscoverySNPCallerPluginV2 was run with the default settings with the following change: the proportion of individuals with the genotype in the locus, the minimum locus coverage, was set to 0.8. Finally, the ProductionSNPCallerPluginV2 was used to convert the data from the local database to VCF format. The Tassel-GBS pipeline does not filter for sequencing depth per se because it is optimised for large numbers of markers in a large sample of individuals at the expense of sequencing depth (optimization of the pipeline is done for sequencing depth of 0.5-3 x) (1). Therefore, the mean sequencing depths in our data ranged between 1.2 – 1310 x for each SNP averaged across individuals. The low coverage in some of the SNPs was compensated by the fact that the minimum locus coverage was set to 0.8 meaning that each SNP was genotyped in at least 80% of the samples. After running the raw data through Discovery and Production pipelines, the resulting number of SNPs was 69,865.

After that, the SNPs were subjected to additional filtering using VCFtools (4). We removed indels, loci with more than two alleles and invariant loci that differed from the reference. However, these invariant sites were retained in the phylogenetic tree construction as they were informative about the divergence from the swan goose. After preliminary analyses we also removed loci with observed heterozygosity over 0.75, because they were potential paralogs mapping to same reference locus. We applied a filter that removed individuals that showed more than 20% missing data across loci. After these filtering steps, we had a dataset that consisted of 33,527 biallelic SNPs and 133 individuals that were successfully genotyped for at least 80% of the loci of which 58 were wild greylags and 75 were domestic geese.

### Population structure analyses

Population clustering and structure at the individual level was analyzed with STRUCTURE 2.3.4 (5) and Principal Component Analysis (PCA) (6, 7) for the whole dataset and within greylags and domestic geese. The Bayesian STRUCTURE approach aims to find an optimal number of clusters (*K*) from a given dataset without prior population/group information by assuming that loci are in linkage equilibrium and each population is in Hardy-Weinberg equilibrium. The clustering of populations is done by considering the individual genotypes and estimating the allele frequencies in populations. For the whole dataset, STRUCTURE was run with 1,000 burn-in steps followed by 10,000 iterations of MCMC for data collection for *K* = 1-10 allowing admixture with five replicates of each run; this appeared to be enough to reach convergence. For the STRUCTURE analyses done separately on greylags and European domestic geese we increased the number of burn-in steps to 10,000 and number of MCMC to 50,000 and set the *K* to 1-7. As with the whole dataset, each run was repeated five times. An admixture model with correlated allele frequencies among populations (8) was used in all the STRUCTURE analyses. The iterations for STRUCTURE analyses were automated with the script StrAuto 1.0 (9). We used both likelihood of *K* and Δ*K* (10) of successive *K* values to determine the optimal number of clusters, as implemented in STRUCTURE HARVESTER (11). CLUMPP 1.1.2 (12) was used to align the assignments from different replicates of *K* and the results were used as an input for visualisation with the program DISTRUCT 1.1 (13). A PCA was performed with prcomp function in R (14) and the significance of the eigenvalues was determined based on the Tracy-Widom distribution (7, 15).

In order to visualize the genetic differences and distance between the reference genome and our data, a neighbor-joining tree was constructed. The tree estimation was performed based on a pairwise distance matrix computed between individuals with the R package ape (16). The *A. cygnoid* reference genome was included in the construction of the neighbor-joining tree and the invariant sites that differed from the reference genome within our data set were also included, thus the tree was constructed with 40,191 loci.

### Tests for admixture and simulations of demographic history

The history of admixture was tested with 3-Population test *f*_3_(C; A, B) implemented in AdmixTools 4.1 (17). This method offers a formal test of admixture that can be used to explain the observed patterns of admixture in a target population and does not require an outgroup. The *f*_3_ test allows separation of ancient polymorphisms from the effects of true admixture, which may be confounded in STRUCTURE. The admixture model is simple, with two source populations contributing to single target population. For identification of admixture between Chinese and European domestics, Grey and White Chinese were combined to represent the Chinese domestic source population and Landes breed that had minimum indication of admixture in STRUCTURE was chosen to represent the European domestic geese source population. The results did not change considerably when other pure breeds were used as a European source (data not shown). Combinations of domestic geese populations and greylag populations were also tested as source populations to see if any of them yielded negative Z-score values supporting history admixture in any of the greylag and domestic goose populations (data not shown).

Different models of demographic history were tested with fastsimcoal2 ver 2.6 (18). Only sites without missing data were used for demographic analyses. We excluded all the SNPs that had missing data within the whole data set and executed the analyses with a site frequency spectrum (SFS) that contained 9,212 polymorphic SNPs. The model estimation utilising the SFS also requires information on the number of monomorphic sites. As there are no estimates of the genetic diversity per base pair for greylags, we made a rough estimation of the proportions of variable and monomorphic sites in our data. The number of bases covered by the GBS tags was calculated from BAM file with –depth option available in SAMtools 1.7 (19). No threshold value was used for this, all the sites that were covered with the tags were recorded, regardless of their sequencing depth or quality. This resulted in 9,801,382 bases covered with tags. After this, we mimicked the filtering steps done for the biallelic SNPs to reduce the total number of sites in equivalent proportions. We removed the same number of sites that corresponded to the number of SNPs that were removed because they were indels, had more than 2 alleles or had heterozygosity over 0.75. Since a proportion of the SNPs were removed from this analysis due to missing data in some of the individuals, we removed an equal proportion of sites from the total number of sites as well. Therefore, the SFS was constructed with 1,681,316 sites in total.

For inferring the demographic history, we chose a subset of individuals from both wild greylags and domestic geese to represent the genetic variation in both groups. This selection was done based on their admixture coefficients from the STRUCTURE analysis so that each wild and domestic population, excluding those that were clearly of a hybrid origin, were represented by an individual with the least amount of admixture from other groups. Therefore, 11 greylags with > 90.8% of greylag ancestry and 15 domestic geese with > 91.4% of European domestic goose ancestry, were selected for the analysis. By doing this, we wanted to minimise the effect of recent admixture on the estimation of divergence time of these two groups, essentially to get the most accurate estimate of the domestication time available. In order to simulate possible population histories, the parameter estimation for each model involved 100,000 simulations and 40 conditional maximization (ECM) cycles. The parameters for each model were estimated with 100 independent runs to obtain the global maximum. The models tested were i) simple divergence of two populations with no migration, ii) divergence of two populations with migration and lastly, iii) divergence of two populations with two different migration matrices (Fig. 2). The best model to represent our data was selected based on Akaike’s weight of evidence as in (18). For parametric bootstrapping 100 SFS were simulated with the parameter estimates obtained from the real SFS, followed by maximum likelihood estimation with 50 independent runs for each bootstrap SFS. The 95% confidence intervals were obtained from the bootstrap data for each estimated parameter.

### The estimation of genetic diversity

Genetic diversity and pairwise *F*_*ST*_ values were investigated using the *hierfstat* R package (20). Expected heterozygosity (H_E_) was calculated for each locus and population and averaged across loci. Difference in average H_E_ between greylags and European domestics was tested with a two-sample t-test with the Welch correction for non-homogeneity of variance (21). For comparing the genetic diversity among wild and domestics, only pure greylag populations (<10% admixture with domestic geese) and pure European domestic geese (<10% admixture with Chinese domestic geese) were used to avoid hybridization effects on the estimates. The admixture proportions were obtained from STRUCTURE analysis detailed above. Therefore, the greylag populations in the Netherlands and Turkey as well as the domestic populations Diepholzer, Crested Faroese, Sebastopol, Toulouse cross, Domestic NY, Buff, Steinbacher and Kholmogor were excluded from the group estimates.

The variance components across loci for hierarchical F statistics for pure greylags and pure European domestics were estimated using hierarchical locus-by-locus analysis of molecular variance (AMOVA, (22)) implemented in Arlequin 3.5.2.1 (23). The significance was tested with 16000 permutations.

## Extended results

### Genetic structure of greylags

Among the wild greylags, the Evanno and likelihood methods indicated that the most likely number of clusters in STRUCTURE was four (Fig. S2). Since the Turkish greylags showed the highest admixture with the domestic geese in the STRUCTURE analysis of the whole data, the STRUCTURE analysis on greylags was executed also without the Turkish greylags but *K* = 4 was deemed best in both analyses. The only difference between the analyses was that without the Turkish greylags, the Danish and Dutch (along with the eastern greylags) showed admixture with a cluster that was not so prominent in the analysis that included the Turkish greylags. The results suggested that there is some population structure, especially within the Iranian populations, but the major separation appears to be between eastern and western greylags. This was also visible in the PCA analysis where there was a tendency to separate the Western European populations from the eastern greylag populations of Iran, Kazakhstan and Russia (Fig. S3). We found three significant PCs (p < 0.05) of which the first PC explained 7.2% of the variation, the second 4.3% and the third 4.1% (Fig. S3). In the neighbor-joining tree, eastern and western greylags mostly fall into separate clades (Fig. S1). Most Dutch and all Danish greylags formed their own clades whereas all Finnish and most of Norwegian greylags were in the same clade (Fig. S1). The Finnish and Norwegian greylags were in different branches of the tree with one individual from Norway being closely related to greylags from SW-Finland. Five samples from an island of Hailuoto in northern Finland formed a subclade that was separate from other Finnish samples that were collected in SW-Finland. The eastern greylags formed a clade with some subclades consistent with the geographical origin of the individuals, but three Norwegian individuals and one Dutch individual also fell into this clade.

### Genetic structure of domestic geese

Based on STRUCTURE analysis conducted solely on domestic geese, the Evanno and likelihood methods indicated that there are two clusters within domestic geese (*K* = 2), approximately representing the European and Chinese domestic geese (Fig. S4). No subsequent split of breeds to different clusters was observed with higher values of *K*. Nearly all European domestic goose populations showed admixture with Chinese domestic geese. The separation between European and Chinese domestic geese was also confirmed by PCA where the first PC separated the Chinese and European domestic geese (Fig. S5). Moreover, within the European domestic geese, the Turkish domestic geese and mostly purebred European domestic geese formed their own groups (Fig. S5). This was also seen in the neighbor-joining tree (Fig. S1). According to Food and Agriculture Organization of the United Nations (FAO), the Diepholzer, Steinbacher and Kholmogor breeds are crosses of the two domestic goose types and they all showed admixture proportions with both types of domestic geese, Diepholzer (88.8% European, 11.2% Chinese), Steinbacher (76.6% European, 23.4% Chinese) and Kholmogor (45.8% European, 54.2% Chinese). In the PCA, Diepholzer and Steinbacher fell into the variation within European domestic goose breeds but Kholmogor was halfway between the European and Chinese domestic geese. Even though the STRUCTURE and PCA did not split breeds into separate genetic clusters, there was a trend of individuals of the same breed forming a clade in the neighbor-joining tree (Fig. S1). The Turkish domestics also formed subclades with individuals from nearby areas with some exceptions.

**Fig. S1.**
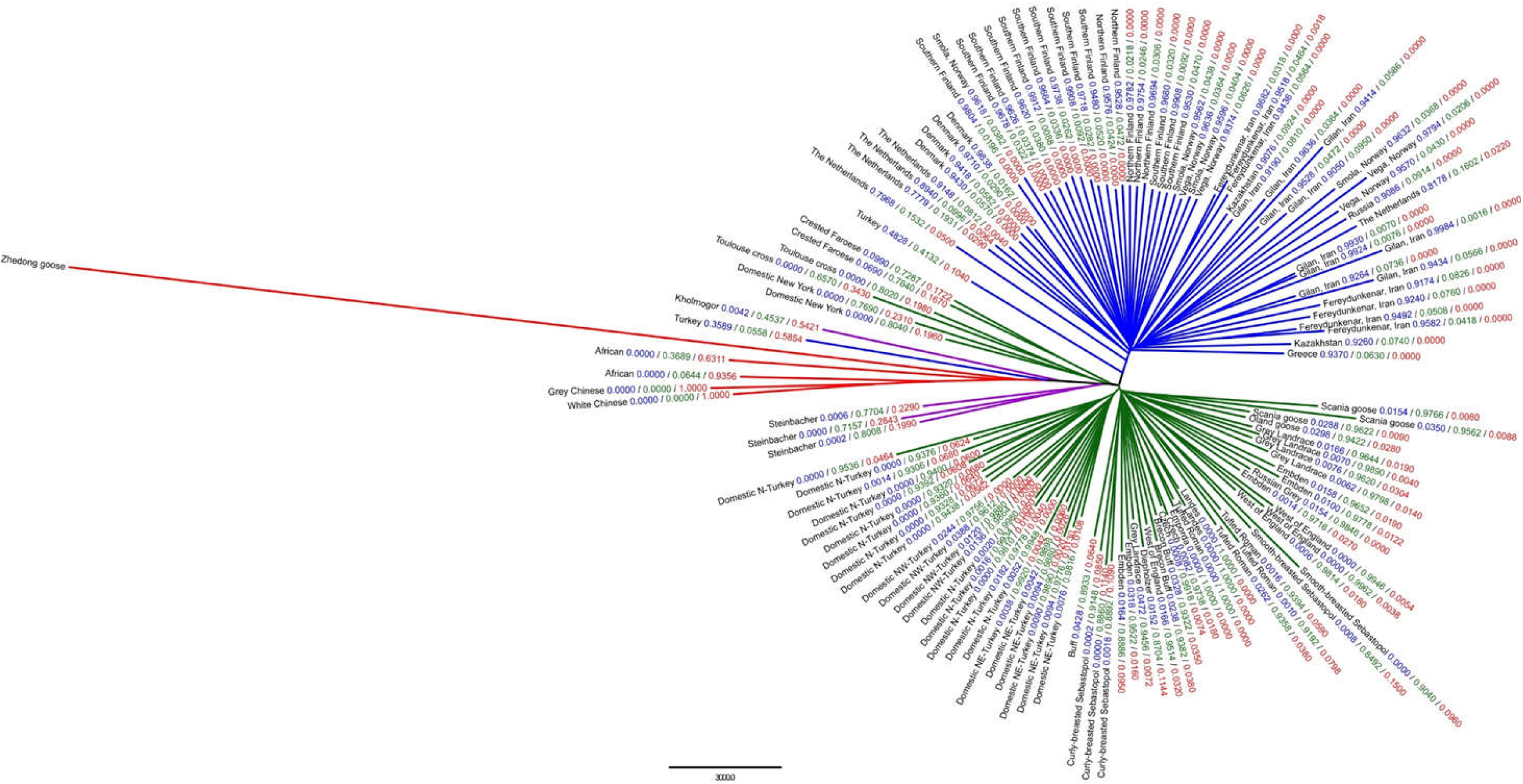
Neighbor-joining tree based on genetic distances between all samples of geese analyzed in this study including the reference genome *A. cygnoid* domesticus breed Zhedong that was used as a reference in SNP calling. Branches leading to greylags are blue, to European domestic geese green and to Chinese domestic geese red. Branches that lead to breeds which are crosses between European and Chinese domestic geese are colored purple. Branches are labelled with a population identifier followed by the greylag, European and Chinese domestic goose admixture proportions from STRUCTURE.

**Fig. S2.**
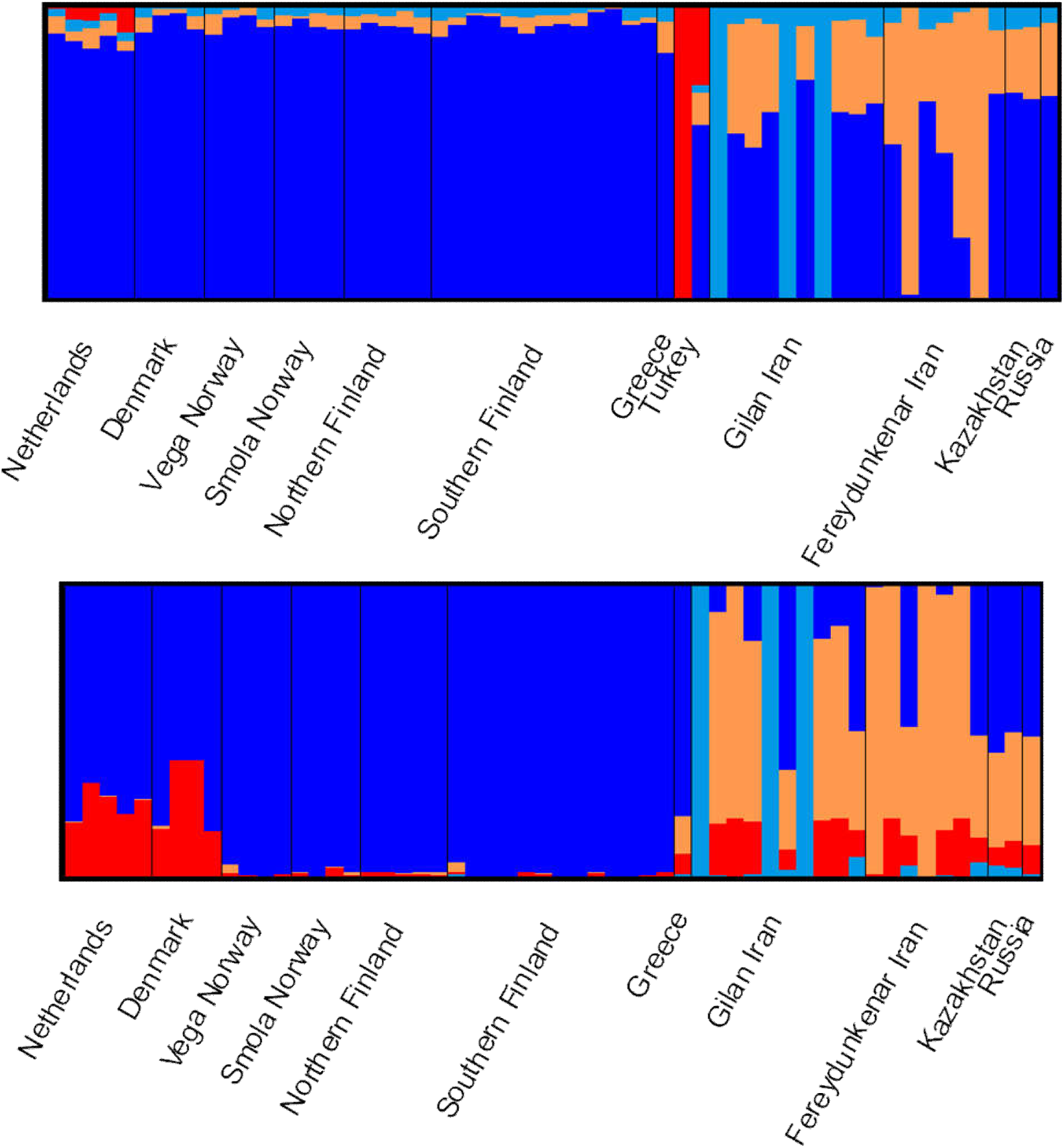
STRUCTURE assignment plots for greylags when *K* = 4. Each vertical line represents one individual with different colors indicating proportion of ancestry from the inferred clusters. The analysis was done with (top) and without (bottom) Turkish greylags because the two Turkish greylags were highly admixed with domestic geese according to STRUCTURE analysis done for the whole data and we wanted to see how that affected the results.

**Fig. S3.**
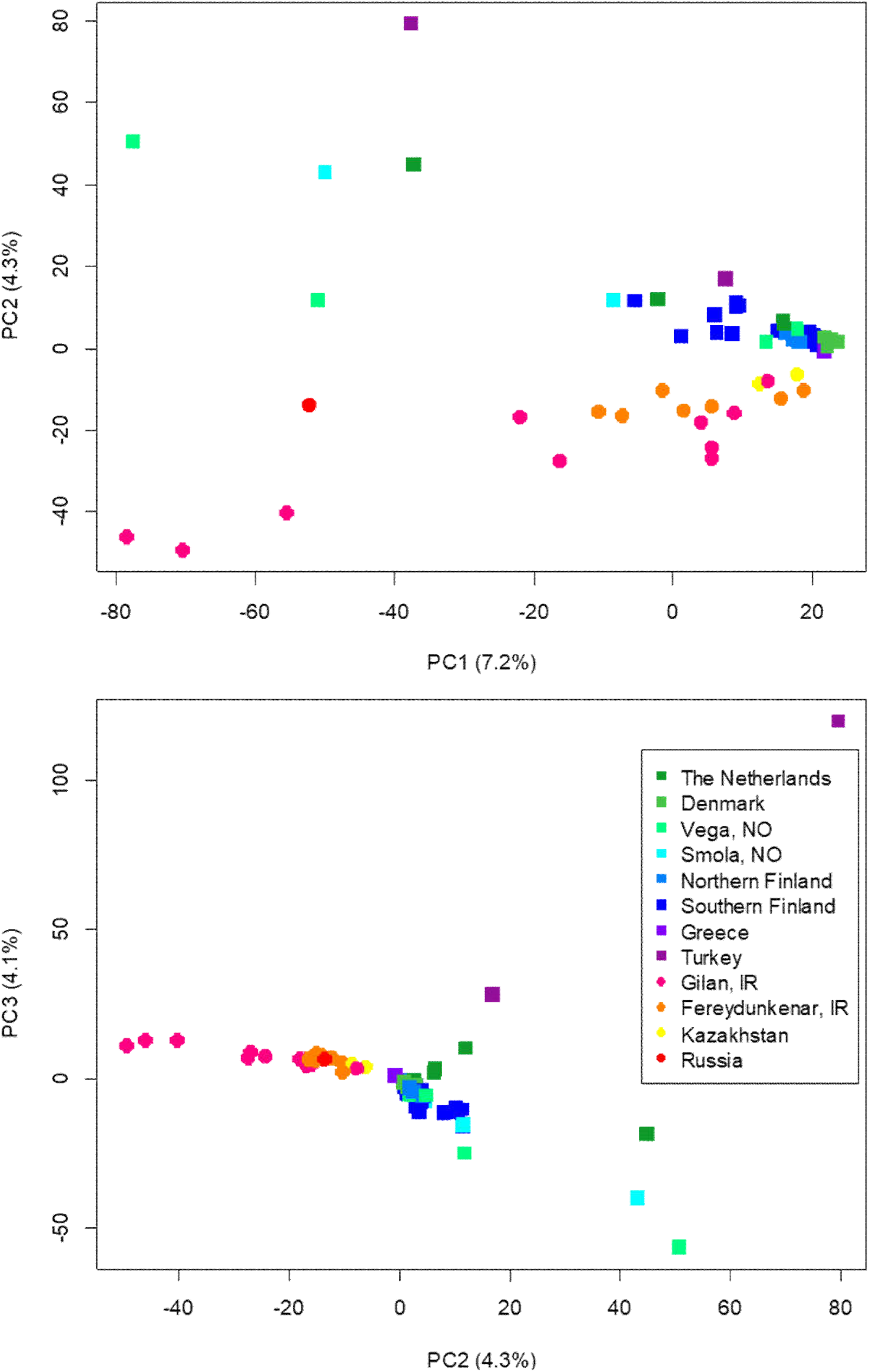
The first three principal components summarizing the genetic variation within wild greylags. European populations are shown with shades of green and blue and purple square symbols while the Asian populations are depicted by yellow, orange and red round symbols. The percentages explained by each PC are shown on the X and Y axes.

**Fig. S4.**
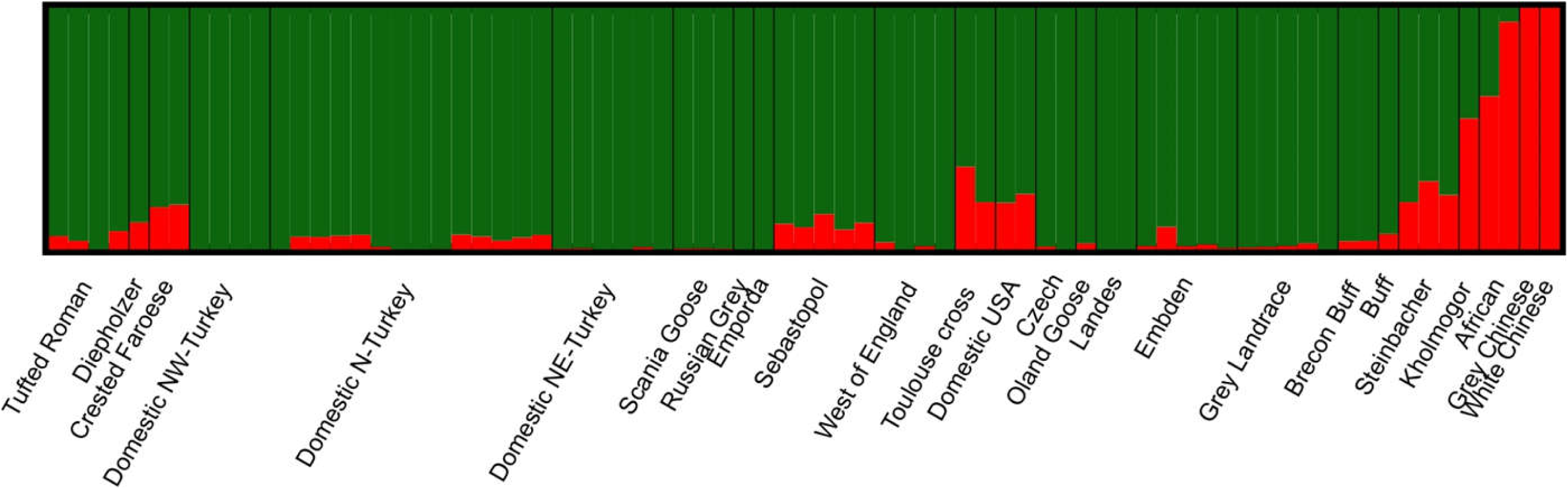
STRUCTURE assignment plot for domestic geese when *K* = 2. Each vertical line represents one individual with different colors indicating proportion of ancestry from the inferred clusters where green indicates proportion of European domestic goose ancestry and red indicates proportion of Chinese domestic goose ancestry in each individual.

**Fig. S5.**
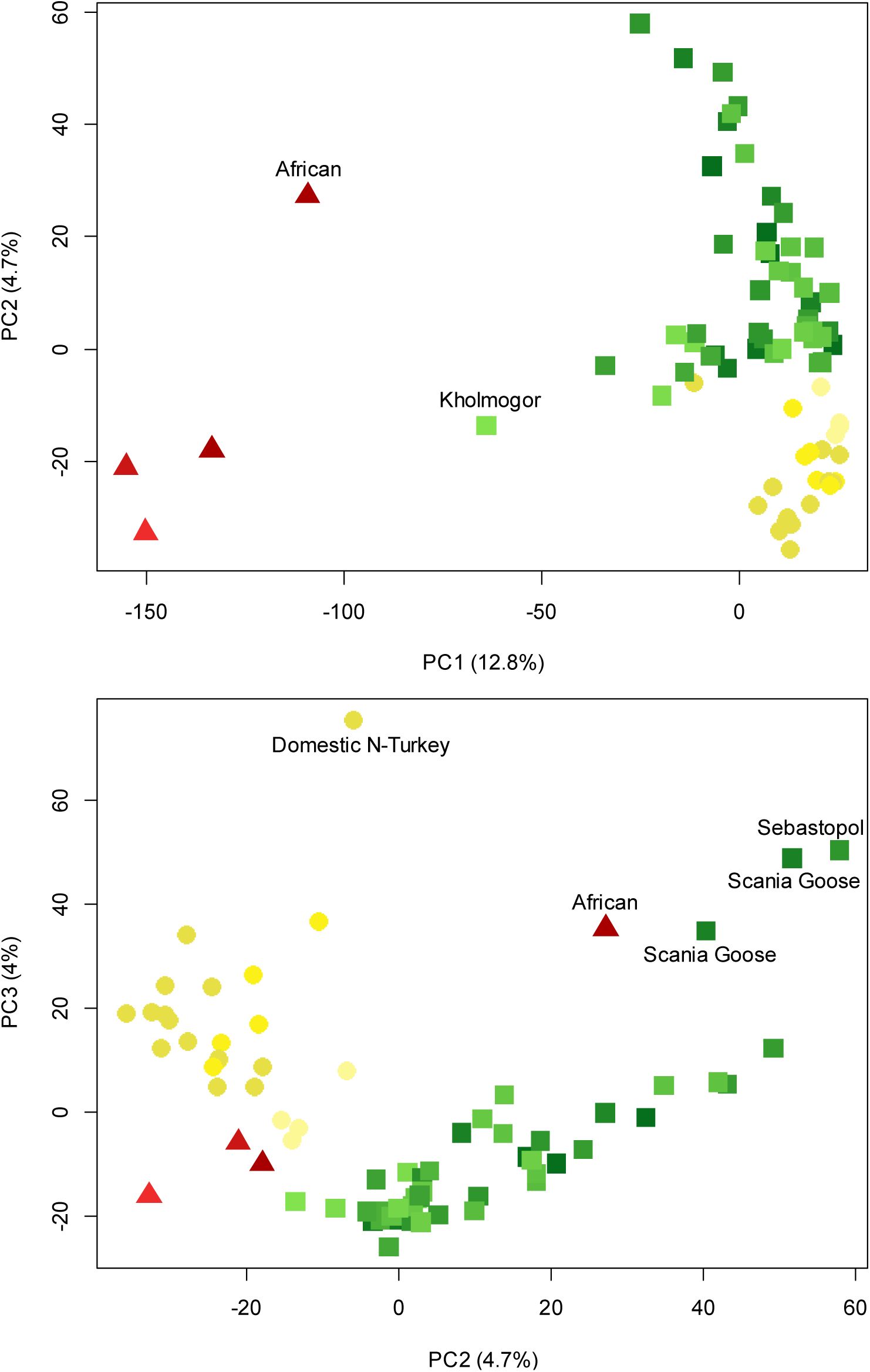
The first three principal components summarizing genetic variation within the domestic geese only. The European breeds are square symbols with different shades of green, Turkish domestics are round symbols with different shades of yellow and Chinese breeds are triangular symbols with different shades of red. The percentages explained by each PC are shown on the X and Y axes.

**Table S1.** Diversity estimates in different greylag populations and breeds of domestic geese. The hybrid status of Diepholzer, Kholmogor and Steinbacher is based on Appendix 1 in Buckland & Guy (2002).

| Population | Status | Sample size | Expected heterozygosity |
| --- | --- | --- | --- |
| Netherlands | Wild Greylag | 5 | 0.157 |
| Denmark | Wild Greylag | 4 | 0.140 |
| Vega, Norway | Wild Greylag | 4 | 0.146 |
| Smola, Norway | Wild Greylag | 4 | 0.148 |
| Northern Finland | Wild Greylag | 5 | 0.147 |
| Southern Finland | Wild Greylag | 13 | 0.149 |
| Greece | Wild Greylag | 1 | NA |
| Turkey | Wild Greylag | 2 | 0.236 |
| Gilan, Iran | Wild Greylag | 10 | 0.145 |
| Fereydunkenar, Iran | Wild Greylag | 7 | 0.142 |
| Kazakhstan | Wild Greylag | 2 | 0.150 |
| Russia | Wild Greylag | 1 | NA |
| Brecon Buff | European Domestic | 2 | 0.110 |
| Buff | European Domestic | 1 | NA |
| Crested Faroese | European Domestic | 2 | 0.117 |
| Czech | European Domestic | 2 | 0.058 |
| Domestic Northern Turkey | European Domestic | 14 | 0.123 |
| Domestic Northeast Turkey | European Domestic | 6 | 0.094 |
| Domestic Northwest Turkey | European Domestic | 4 | 0.092 |
| Domestic NY | European Domestic | 2 | 0.143 |
| Embden | European Domestic | 5 | 0.120 |
| Emporda | European Domestic | 1 | NA |
| Grey Landrace | European Domestic | 5 | 0.107 |
| Landes | European Domestic | 2 | 0.047 |
| Oland Goose | European Domestic | 1 | NA |
| Russian Grey | European Domestic | 1 | NA |
| Scania Goose | European Domestic | 3 | 0.099 |
| Sebastopol | European Domestic | 5 | 0.133 |
| Toulouse cross | European Domestic | 2 | 0.151 |
| Tufted Roman | European Domestic | 4 | 0.113 |
| West of England | European Domestic | 4 | 0.093 |
| Diepholzer | European x Chinese Domestic | 1 | NA |
| Steinbacher | European x Chinese Domestic | 3 | 0.152 |
| Kholmogor | European x Chinese Domestic | 1 | NA |
| African | Chinese Domestic | 2 | 0.224 |
| Grey Chinese | Chinese Domestic | 1 | NA |
| White Chinese | Chinese Domestic | 1 | NA |

**Table S2.**
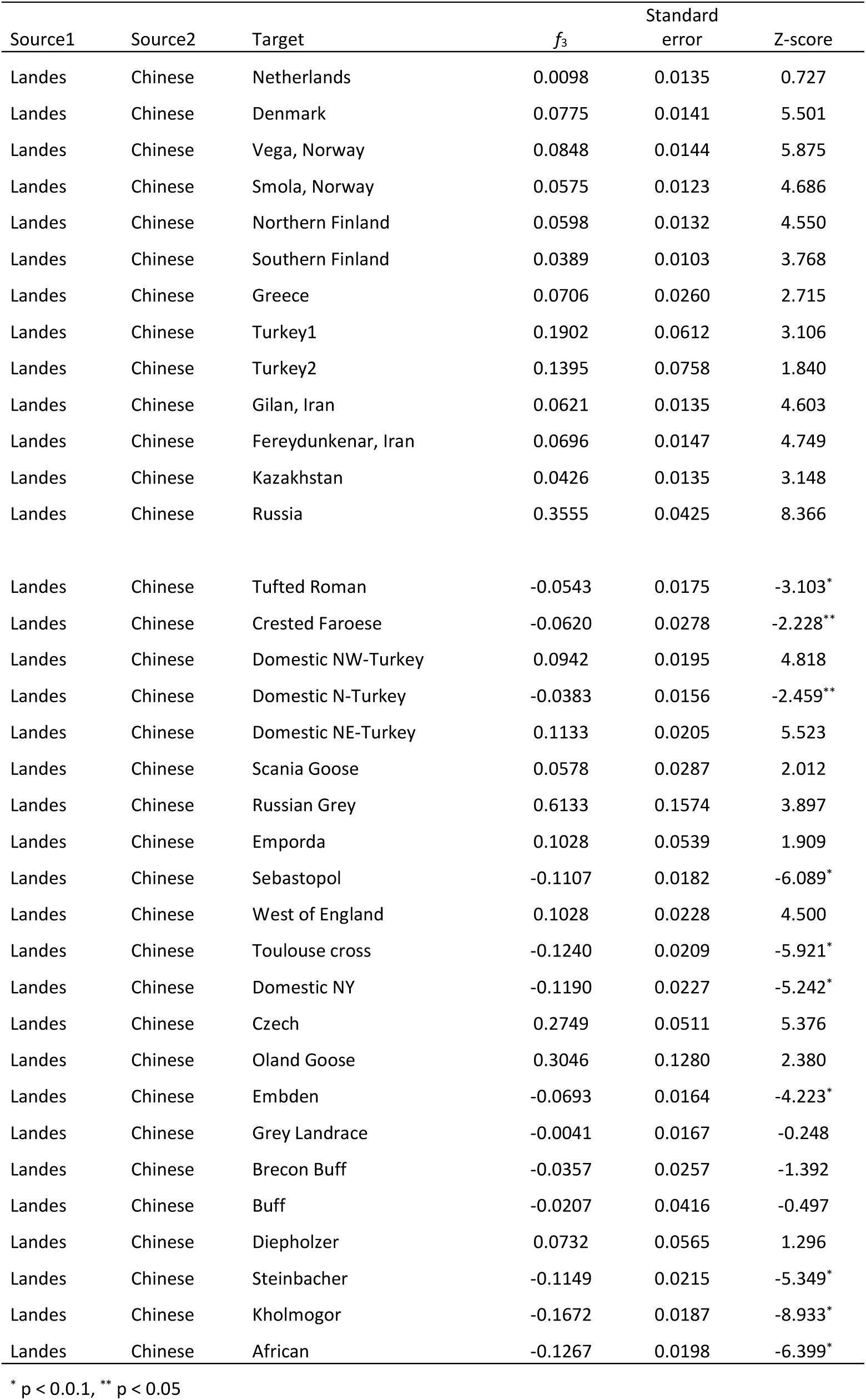
The Patterson’s 3-Population test statistics obtained to test the history of admixture in different greylag populations and breeds of domestic geese.

**Table S3.** Model selection results and parameter estimates for different demographic models that were tested (see text). Confidence intervals for the best model are shown below the bottom line.

| Model | Number of parameters | log L | AIC | ΔAIC | AIC<sub>W</sub> | ANCSIZE | N<sub>DOM</sub> | N<sub>WILD</sub> | T<sub>1</sub> | T<sub>2</sub> | M1<sub>WD</sub> | M1<sub>DW</sub> | M2<sub>WD</sub> | M2<sub>DW</sub> |
| --- | --- | --- | --- | --- | --- | --- | --- | --- | --- | --- | --- | --- | --- | --- |
| Divergence with 2 migration matrices | 9 | -29766.4 | 59550.7 | 0 | 0.99999999 | 1112 | 959 | 2504 | 5319 | 159 | 5.35x10<sup>-4</sup> | 4.25x10<sup>-4</sup> | 6.69x10<sup>-4</sup> | 1.72x10<sup>-3</sup> |
| Divergence with migration | 6 | -29787.7 | 59587.44 | 36.736 | 1.0541E-08 | 756 | 1056 | 2304 | 5952 |  | 7.70x10<sup>-4</sup> | 8.86x10<sup>-4</sup> |  |  |
| Divergence without migration | 4 | -29919.8 | 59847.53 | 296.83 | 3.5009E-65 | 7312 | 1047 | 2576 | 226 |  |  |  |  |  |
|  |  |  |  |  |  | 291.95-9030.75 | 879.95-1073.25 | 2341.60-2695.55 | 1538.80-8225.45 | 79.65-462.95 | 2.96x10<sup>-4</sup>-6.75x10<sup>-4</sup> | 1.73x10<sup>-7</sup>-6.03x10<sup>-4</sup> | 4.57x10<sup>-4</sup>-9.69x10<sup>-4</sup> | 1.33x10<sup>-3</sup>-2.15x10<sup>-3</sup> |

ANCSIZE: effective population size of ancestral population, N<sub>DOM</sub>: effective population size for domestic geese, N<sub>WILD</sub>: effective population size for greylags, T<sub>1</sub>: time of divergence in generations, T<sub>2</sub>: estimate of time in generations when the migration matrix switched, M1<sub>WD</sub>: migration rate from wild to domestic following T1, M1<sub>DW</sub>: migration rate from domestic to wild following T1, M2<sub>WD</sub>: migration rate from wild to domestic following T2, M2<sub>DW</sub>: migration rate from domestic to wild following T2

**Table S4. (separate file)** Pairwise *F*_*ST*_ values for each population analysed in this study.

